# Did viruses evolve as a distinct supergroup from common ancestors of cells?

**DOI:** 10.1101/049171

**Authors:** Ajith Harish, Aare Abroi, Julian Gough, Charles Kurland

**Affiliations:** Structural and Molecular Biology Group, Department of Cell and Molecular Biology, Biomedical Center, Uppsala University, 751 24 Uppsala, Sweden.; Estonian Biocentre, Riia 23, Tartu, 51010, Estonia.; Computational Genomics Group, Department of Computer Science, University of Bristol, The Merchant Venturers Building, Bristol BS8 1UB, UK; Microbial Ecology, Department of Biology, Lund University, Ecology Building, SE-223 62 Lund, Sweden.

## Abstract

The evolutionary origins of viruses according to marker gene phylogenies, as well as their relationships to the ancestors of host cells remains unclear. In a recent article Nasir and Caetano-Anollés reported that their genome-scale phylogenetic analyses identify an ancient origin of the “viral supergroup” (Nasir et al (2015) A phylogenomic data-driven exploration of viral origins and evolution. *Science Advances*, 1(8):e1500527). It suggests that viruses and host cells evolved independently from a universal common ancestor. Examination of their data and phylogenetic methods indicates that systematic errors likely affected the results. Reanalysis of the data with additional tests shows that small-genome attraction artifacts distort their phylogenomic analyses. These new results indicate that their suggestion of a distinct ancestry of the viral supergroup is not well supported by the evidence.

## Introduction

The debate on the ancestry of viruses is still undecided: In particular, it is still unclear whether viruses evolved before their host cells or if they evolved more recently from the host cells. The virus-early hypothesis posits that viruses predate or coevolved with their cellular hosts [1]. Two alternatives describe the virus-late scenario: (i) the progressive evolution also known as the escape hypothesis and (ii) regressive evolution or reduction hypothesis. Both propose that viruses evolved from their host cells [1]. According to the first of these two virus-late models, viruses evolved from their host cells through gradual acquisition of genetic structures. The other alternative suggests that viruses, like host-dependent endoparasitic bacteria, evolved from free-living ancestors by reductive evolution. The recent discovery of the so-called giant viruses with double-stranded DNA genomes that parallel endoparasitic bacteria with regards to genome size, gene content and particle size revived the reductive evolution hypothesis. However, there are so far no identifiable ‘universal’ viral genes that are common to viruses such as the ubiquitous cellular genes. In other words, examples of common viral components that are analogous to the ribosomal RNA and ribosomal protein genes, which are common to cellular genomes, are not found. This is one compelling reason that phylogenetic tests of the “common viral ancestor” hypotheses seem so far inconclusive.

Recently, Nasir and Caetano-Anollés [2] employed phylogenetic analysis of whole-genomes and gene contents of thousands of viruses and cellular organisms to test the alternative hypotheses. The authors conclude that viruses are an ancient lineage that diverged independently and in parallel with their cellular hosts from a universal common ancestor (UCA). They reiterate their earlier claim [3] that viruses are a unique lineage, which predated or coevolved with the last UCA of cellular lineages (LUCA) through reductive evolution rather than through more recent multiple origins. Their claims are based on analyses of statistical-and phyletic distribution patterns of protein domains, classified as superfamilies (SFs) in Structural Classification of Proteins (SCOP) [4].

Detailed re-examination of Nasir and Caetano-Anollés’ phylogenomic approach [2, 3] suggests that small genomes systematically distort their phylogenetic reconstructions of the tree of life (ToL), especially the rooting of trees. Here the ToL is described as the evolutionary history of contemporary genomes: as a tree of genomes or a tree of proteomes (ToP) [5-8]. The bias due to highly reduced genomes of parasites and endosymbionts in genome-scale phylogenies has been known for over a decade [5, 6]. In fact, prior to the recent proposal [2] these authors recognized the anomalous effects of including small genomes in reconstructing the ToL in analyses that were limited to cellular organisms [9] or which included giant viruses [3]: As they say “In order to improve ToP reconstructions, we manually studied the lifestyles of cellular organisms in the total dataset and excluded organisms exhibiting parasitic (P) and obligate parasitic (OP) lifestyles, as their inclusion is known to affect the topology of the phylogenetic tree” [3]. But, they may not have adequately addressed this problem, particularly when the samplings include viral genomes that are likely to further exacerbate bias due to small genomes [2, 3]. For this reason we systematically tested the reliability of the phylogenetic trees, especially the rooting approach favored by Nasir and Caetano-Anollés [2]. This approach depends critically on a pseudo-outgroup to root the ToP (ToL), but that pseudo-outgroup is not identified empirically. Rather, it is assumed *a priori* to be an empty set [2].

We show here in several independent phylogenetic reconstructions that a rooting based on a hypothetical “all-zero” outgroup—an ancestor that is assumed to be an empty set of protein domains—creates specific phylogenetic artifacts: In this particular approach [2, 3] implementing the all-zero outgroup artifactually draws the taxa with the smallest genomes (proteomes) into a false rooting very much like the classical distortions due to long-branch attractions (LBA) in gene trees [10].

## Results and Discussion

We emphasize at the outset of this study that virtually all the evolutionary interpretations based on phylogenetic reconstructions depend on a reliable identification of the root of a tree [11, 12]. In particular, the rooting of a tree will determine the branching order of species, and define the ancestor-descendant relationships between taxa as well as the derived features of characters (character states). In effect, the root polarizes the order of evolutionary changes along the tree with respect to time. By the same token, the rooting of a tree distinguishes ancestral states from derived states among the different observed states of a character. If ancestral states are identified directly, for example from fossils, characters can be polarized *a priori* with regard to determining the tree topology. *A priori* polarization provides for intrinsic rooting. However, direct identification of ancestral states, particularly for extant genetic data is rarely possible. Consequently conventional phylogenetic methods use time-reversible (undirected) models of character evolution and they only compute unrooted trees. For example, the root of the iconic ribosomal RNA ToL or any other gene-tree has not been determined directly [13, 14].

Accordingly, conventional approaches to character polarization are indirect and the rooting of trees is normally a two-stage process. The most common rooting method is the outgroup comparison method, which is based on the premise that character-states common to the ingroup (study group) and a closely related sister-group (the outgroup) are likely to be ancestral to both. Therefore, in an unrooted tree the root is expected to be positioned on the branch that connects the outgroup to the ingroup. In this way, the tree (and characters) may be polarized *a posteriori* [12, 14]. However, there are no known outgroups for the ToL. In the absence of natural outgroups, pseudo-outgroups are used to root the ToL [14]. The best-known case is the root grafted onto the unrooted ribosomal RNA ToL based on presumed ancient (pre-LUCA) gene duplications [15, 16]. Here the paralogous proteins act as reciprocal outgroups that root each other. Unlike gene duplications used to root gene trees, the challenge of identifying suitable outgroups becomes more acute for genome trees.

To meet this challenge Nasir and Caetano-Anollés use a hypothetical pseudo-outgroup: an artificial taxon constructed from presumed ancestral states. For each character (SF), the state ‘0’ or ‘absence’ of a SF is assumed to be “ancestral” *a priori*. This artificial “all-zero” taxon is used as outgroup to root the ToL. Further, they use the Lundberg rooting method, in which outgroups are not included in the initial tree reconstruction. The Lundberg method involves estimating an unrooted tree for the ingroup taxa only, and then attaching outgroup(s) (when available) or a hypothesized ancestor to the tree *a posteriori* to determine the position of the root [17]. Unrooted trees describe relatedness of taxa based on graded compositional similarities of characters (and states). Accordingly, we can expect the “all-zero” pseudo-outgroup to cluster with genomes (proteomes) in which the smallest number of SFs is present. The latter are the proteomes described by the largest number of ‘0s’ in the data matrix.

The instability of rooting with an all-zero pseudo-outgroup becomes clear when the smallest proteome in a given taxon sampling varies in the rooting experiments (Figs. 1 and 2). Rooting experiments were preformed both for SF occurrence (presence/absence) patterns and for SF abundance (copy number) patterns. However, we present results for the SF abundance patterns, as in [2]. Throughout, we refer to genomic protein repertoires as proteomes. Proteome size related as the number of distinct SFs in a proteome (SF occurrence) is depicted next to each taxon for easy comparison in Figs. 1 and 2. Phylogenetic analyses were carried out as described in [2, 3] (see methods). SFs that are shared between proteomes of viruses and cellular organisms were used as the characters (Fig. 1a) as in [2]. Initially no viruses are included in tree reconstructions and here the root was placed within the Archaea, which has the smallest proteome (503 SFs) among the supergroups (Fig 1b). When a still smaller bacterial proteome (420 SFs) was included, the position of the root as well as the branching order changed. In this case, the bacteria were split into two groups and the root was placed within one of the bacterial groups (Fig. 1c). Further, when a much smaller archaeal proteome was included (228 SFs), the root was relocated to a branch leading to the now smallest proteome (Fig. 1d). Note that the newly included taxa, both bacteria and archaea are host-dependent symbionts with reduced genomes.

**Figure 1.**
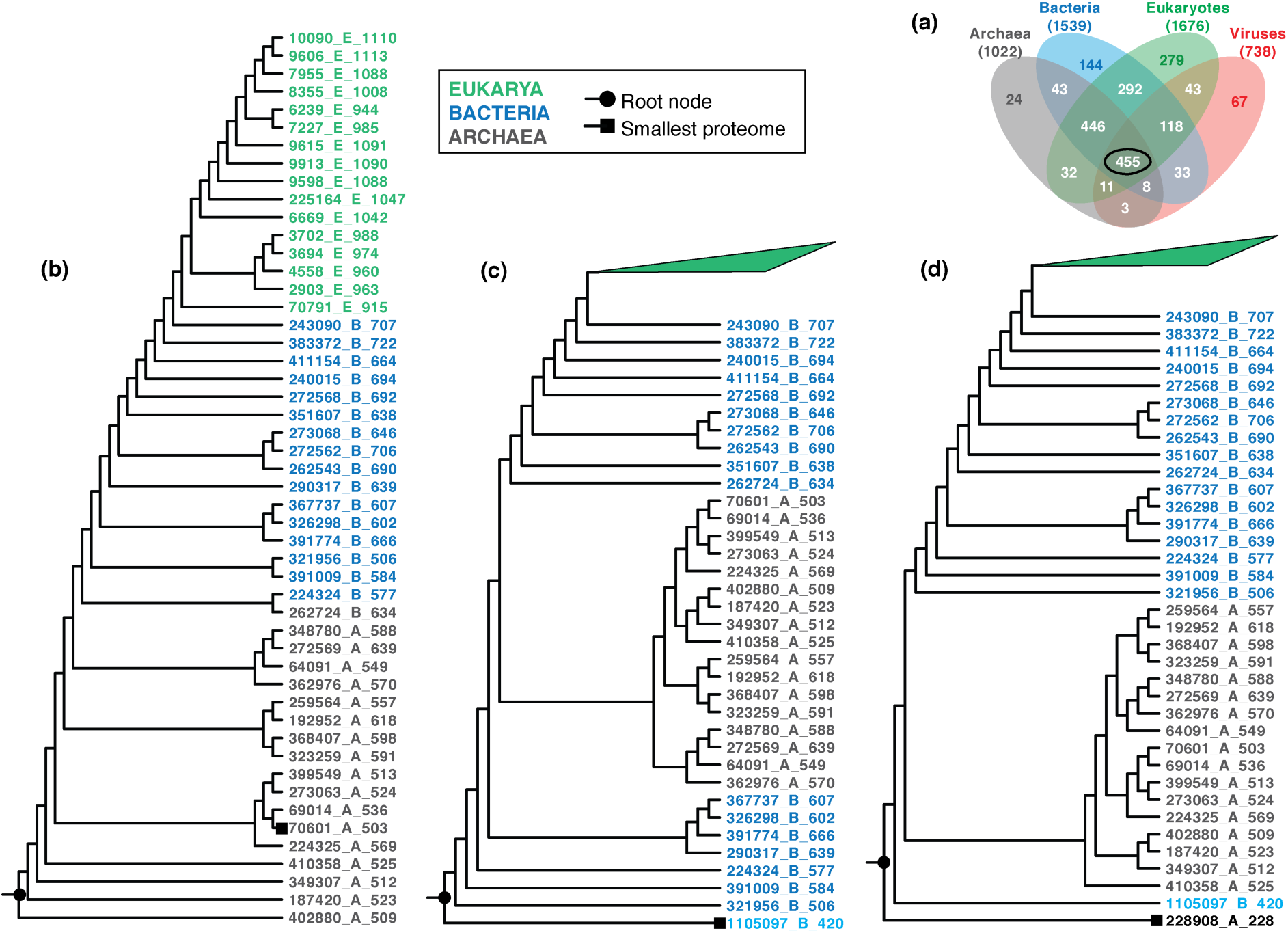
Implementing an “all-zero” pseudo-outgroup [2] severely distorts rooting of the ToL. Rooted trees were reconstructed from a subset of 368 taxa (proteomes) sampled in [2], which included 17 taxa each from Archaea, Bacteria, Eukarya (ABE) and 9 taxa from the virus groups (V). (a) Venn diagram shows the 455 SFs shared between viruses and cells (ABEV), which were used to reconstruct trees. (b) Single most parsimonious tree of ABE taxa rooted within Archaea. (c, d) New taxa, which represent the smallest proteome after inclusion, were progressively included in size order. The position of the root node changed accordingly to the branch corresponding to a group (or taxon) with the smallest proteome, which is Bacteria (c), Archaea (d); the Eukarya section is collapsed since tree topology is unaffected. Taxa are described by their NCBI taxonomy ID, taxonomic affiliation (A, B, E or V) and proteome size in terms of the number of distinct SFs present in the genome. To compare the position of the root node trees are drawn to show branching patterns only, branch lengths are not proportional to the quantity of evolutionary change.

**Figure 2.**
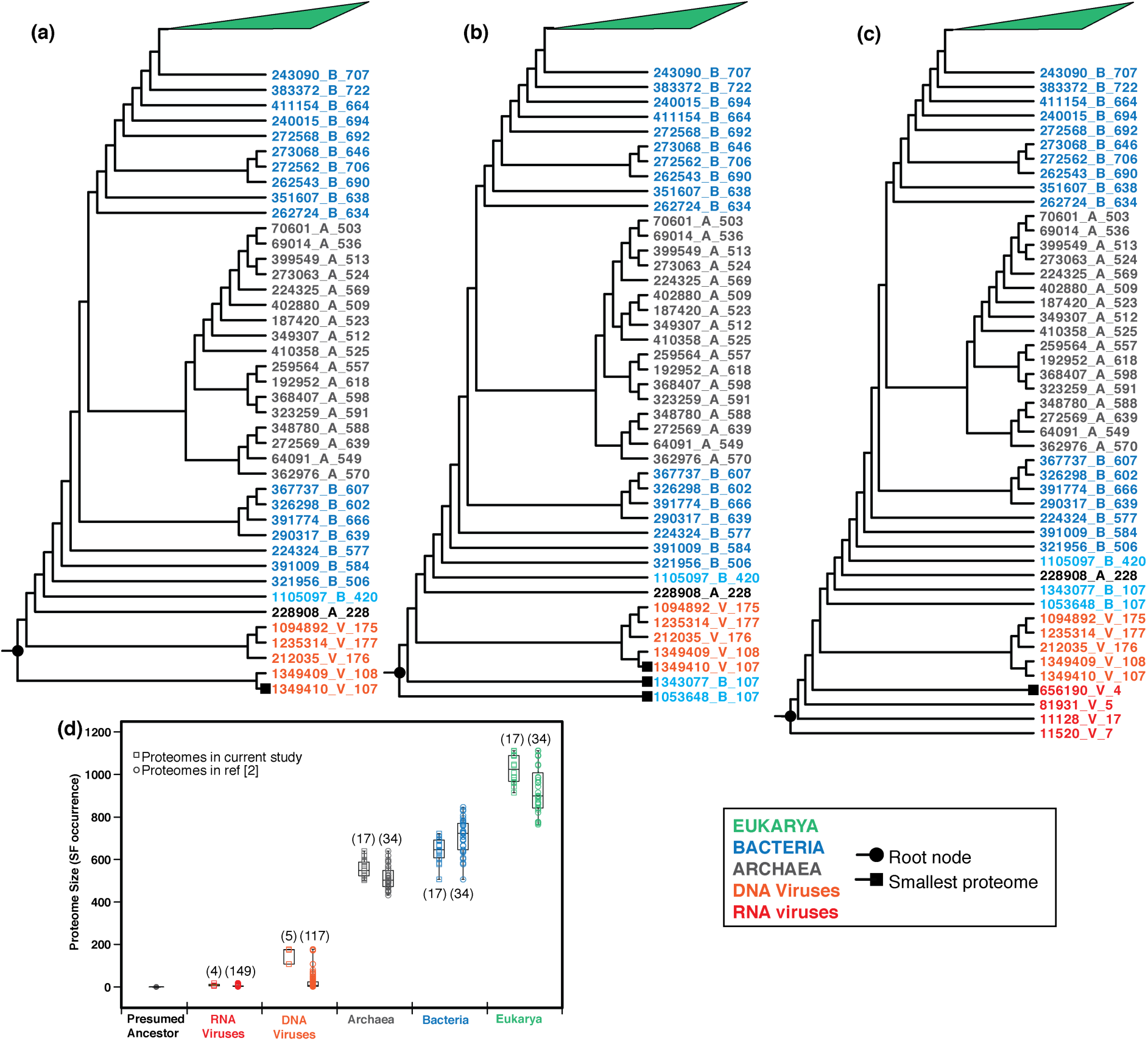
Rooting experiments (continued from Fig. 1) show rooting bias of “all-zero” pseudo-outgroup towards small proteomes. (a-c) New taxa, which represent the smallest proteome after inclusion, were progressively included in size order, where the smallest proteome was from mega DNA viruses (a), Bacteria (b) and RNA viruses (c). Details in trees are same as in Fig. 1. (d) Comparison of proteome sizes of sampled taxa in terms of SF occurrence used to estimate trees in this study and in [2]. Numbers in parentheses above each group indicate the number of proteomes.

Similarly including viruses in the analyses draws the root towards the smaller viral proteomes (Fig. 2). As in the rooting experiments in Fig. 1, a group of DNA viruses (107-175 SFs) was introduced. These DNA viruses have larger proteomes than do the RNA viruses, but they are much smaller than most known endosymbiotic bacteria (Fig. 2d). Again, the root was repositioned within the DNA viruses group (Fig. 2a). Following this experiment, two extremely reduced (107 SFs each) endosymbiotic bacteria classified as Betaproteobacteria were included. These further displaced the root closer to the smallest set of proteomes (Fig. 2b). Finally, a set of four RNA viruses (4-17 SFs) in their genomes was introduced and they rooted the tree within the RNA viruses (Fig. 2c). These results challenge the conclusion drawn previously that the proteomes of RNA viruses are more ancient than proteomes of DNA viruses [2]. In addition, the results contradict the purported antiquity of viral proteomes as such. Rather, the data suggest that there are severe artifacts generated by genome size-bias due to the inclusion of the viral proteomes in the analysis. These artifacts are expressed as grossly erroneous rootings caused by small-genome attraction (SGA) in the Lundberg rooting using the hypothetical all-zero outgroup.

Including the all-zero pseudo-outgroup in the analysis either implicitly (defined by the ANCSTATES option in PAUP*) or explicitly (as a taxon in the data matrix) does not make a difference to the tree topology and rooting. We note that including the hypothetical ancestor during tree estimation amounts to *a priori* character polarization and pre-specification of the root. In addition, the position of the root was the same in the different rooting experiments when the all-zero pseudo-ancestor was explicitly specified as the outgroup for root trees using the outgroup method (see supplementary Figs S1 & S2). These rooting experiments reveal a strong bias in rooting that favors small genomes irrespective of the different rooting methods—outgroup rooting, Lundberg rooting and intrinsic rooting—used to root the ToL with the “all-zero” hypothetical ancestor. This SGA artifact is comparable to the better-known LBA artifact that is associated with compositional bias of nucleotides or of amino acids that distort gene trees [10].

The use of artificial outgroups is not uncommon in rooting experiments when rooting is ambiguous [11]. Artificial taxa are either an all-zero outgroup or an outgroup constructed by randomizing characters and/or character-states of real taxa. Although rooting experiments with multiple real outgroups, or randomized artificial outgroups that simulate loss of phylogenetic signal can minimize the ambiguity in rooting the all-zero outgroup has proved to be of little use [11, 14]. Conclusions based on an all-zero outgroup are often refuted when empirically grounded analysis with real taxa are carried out [14]. Indeed, the present rooting experiments (Figs. 1 and 2) clearly show that the position of the root depends on the smallest genome in the sample when a hypothetical all-zero ancestor/outgroup is used. In effect, the rooting approach favored by Nasir and Caetano-Anolles [2] is not reliable.

Nevertheless, small proteome size is not an irreconcilable feature of genome-tree reconstructions [5, 6, 18]. Small genome attraction artifacts may be observed when highly reduced proteomes of obligate endosymbionts are included in analyses with common samplings [3, 5, 6, 18]. However, their untoward effects are only observed in the shallow end of the tree where the endosymbionts might be clustered with unrelated groups. In such cases the deep divergences or clustering of major branches are unperturbed [6, 18]. Such size-biases are readily corrected by normalizations that account for genome size (actually specific SF content in this case) [5, 6, 18].

Though size-bias correction for SF content phylogeny is known to be reliable, both for measures based on distance [6] and those based on frequencies of character distribution [18], Nasir and Caetano-Anollés do not apply such corrections. They only attend to the proteome size variations associated with SF abundances [2, 3]. Instead of accounting for novel taxon-specific SFs in their model of evolution, the authors choose to exclude potentially problematic small proteomes of parasitic bacteria and to include only the proteomes of ‘free-living’ cellular organisms in their analyses [2, 3]. But all viruses are parasites and obviously even more extreme examples of minimal proteomes.

This problem is further exacerbated by the uneven and largely incomplete annotation of SF domains in viral proteins [19, 20]. In fact, many viral ‘proteomes’ that were sampled in [2] are as small as a single SF. It is not clear why the inclusion of small viral proteomes was not recognized as even more problematic than the inclusion of small parasitic bacterial proteomes, in spite of the previous assertion of these authors that small proteomes should be excluded [9]. Nevertheless, including small viral proteomes is inconsistent with specifically excluding small cellular proteomes in the ToL, especially when hypotheses of reductive evolution are considered. Screening taxa based on ‘lifestyle’ (free-living or parasitic) seems unwarranted since extreme reductive genome evolution, sometimes called genome streamlining, is not limited to host adapted parasitic bacteria but is common in free-living bacteria as well as eukaryotes [21-23].

In addition to the ToP, the authors use a so-called tree of domains (ToD) to support their conclusion that proteomes of viruses are ancient and that proteomes of RNA viruses are particuarly ancient. The ToD is projected as the evolutionary trajectory of individual SFs. Such projections are used as proxies to determine the relative antiquity or novelty of SFs [2]. The ToD like the ToP is also rooted with a presumed pseudo-outgroup and that rooting may be an artifact as for the ToP. Much more serious than potential artifacts in ToD is an egregiously bad assumption from the perspective of the SCOP hierarchal classification: the very notion that the ToD describes evolutionary relationships between SFs in the same way the ToL describes genealogy of species. Evolutionary relationships between SFs with different folds in SCOP classification are not established [4, 24]. Physicochemical protein folding experiments and corresponding statistical analyses of sequence evolution patterns, including simulations of protein folding are all consistent with the observations that the sequence-structure space of SFs is discontinuous [25, 26]. Empirical data indicate that the evolutionary transition from one SF to another through gradual changes implied in the ToD is unlikely, if at all feasible. This makes the ToD hypothesis, which assumes that all SFs are related to one another by common ancestry untenable. Thus the ToD contradicts the very basis upon which SFs are classified in the SCOP hierarchy [4, 24]. The ToD is therefore uninterpretable as an evolutionary history of individual SFs. Accordingly, the ToD cannot reflect the ‘relative ages’ of SFs nor can it support the inferred antiquity of viruses in the ToP.

Unlike phylogenetic trees that describe the evolution of individual proteomes, Venn diagrams, SF sharing patterns and summary statistics of SF frequencies among groups of proteomes only depict generalized trends. Multiple evolutionary scenarios can be invoked *a priori* to explain the general trends without any phylogenetic analyses. Although such patterns may be suggestive, they do not by themselves support reliable phylogenetic inferences [19, 20]. Thus the authors’ inferences in [2] based on statistical distributions of SFs alone are speculative, at best.

In summary, the authors’ [2, 3] proposed rooting for the ToL seems to be influenced by clearly identifiable artifacts. The conceptual issue of proteome evolution may be traced to a very widely held view: namely that “ToP were rooted by the minimum character state, assuming that modern proteomes evolved from a relatively simpler urancestral organism that harbored only few FSFs” [2, 9]. Thus, a common prejudice is that the origins of modern proteomes ought to reflect a monotonic evolutionary progression that is embodied in something like Aristotle’s Great Chain of Being [10]. The “all-zero” or “all-absent” hypothetical ancestor is neither empirically grounded nor biologically meaningful, but it does indeed reflect a common prejudice [10, 14]. Indeed, we bring into question the inferred relative antiquity of viruses and Archaea in the ToL and the notion that viruses make up an independent fourth supergroup [2]. In effect, we suggest that the phylogenetic approach of Nasir and Caetano-Anolles’ [2, 3] provides neither a test nor a confirmation of any one of the hypotheses for the origins of viruses [1]. Despite its importance, reconciling the extensive genetic and morphological diversity of viruses as well as their evolutionary origins remains murky [1, 27]. Better methods and empirical models are required to test whether a multiplicity of scenarios or a single over-arching hypothesis is applicable to understand the origins of viruses.

## Methods

Here, genomic protein repertoires are referred to as proteomes. We re-analyzed a subset of the 368 proteomes sampled in [2] for phylogenetic rooting of diverse cells and their viruses. Here, we sampled 102 cellular proteomes containing 34 each from Archaea, Bacteria and Eukaryotes, respectively as well as 16 viral proteomes from [2]. For the latter, we note that the DNA virus proteomes were substantially larger than those of RNA viruses in terms of the number of identified SFs. In addition we included for comparison some of the smallest known proteomes of Archaea and Bacteria not included in [2]. Roughly, half of the sampled proteomes were analyzed (Figs 1 and 2) for computational simplicity. Results did not vary when all the sampled taxa were included (see supplementary Figs S3 & S4). Rooting experiments were preformed both with SF occurrence (presence/absence) patterns and SF abundance (copy number) patterns; however, we present results for the SF abundance patterns as in [2]. In addition to the Lundberg rooting procedures carried out as in [2, 3], rooting experiments were repeated by including the all-zero taxon in the tree reconstruction process implicitly (using the ANCSTATES option in PAUP*) and explicitly as taxon in the data matrix. Further, when the all-zero taxon was explicitly included, rooting experiments were also repeated with the outgroup rooting method. Phylogenetic reconstructions were carried out using maximum parsimony criteria implemented in PAUP* ver. 4.0b10 [28] with heuristic tree searches using 1,000 replicates of random taxon addition and tree bisection reconnection (TBR) branch swapping. Trees were rooted by Lundberg method, outgroup method or intrinsically rooted by including the hypothetical all-zero ancestor in tree searches.

## Acknowledgements

A. H. acknowledges support from The Swedish Research Council (to Måns Ehrenberg) and the Knut and Alice Wallenberg Foundation, RiboCORE (to Måns Ehrenberg), A.A. acknowledges support from European Regional Development Fund through the Centre of Excellence in Chemical Biology grant number 3.2.0101.08-0017 and C.G.K. acknowledges support from the Nobel Committee for Chemistry of the Royal Swedish Academy of Sciences.

---

**Figure S1.**
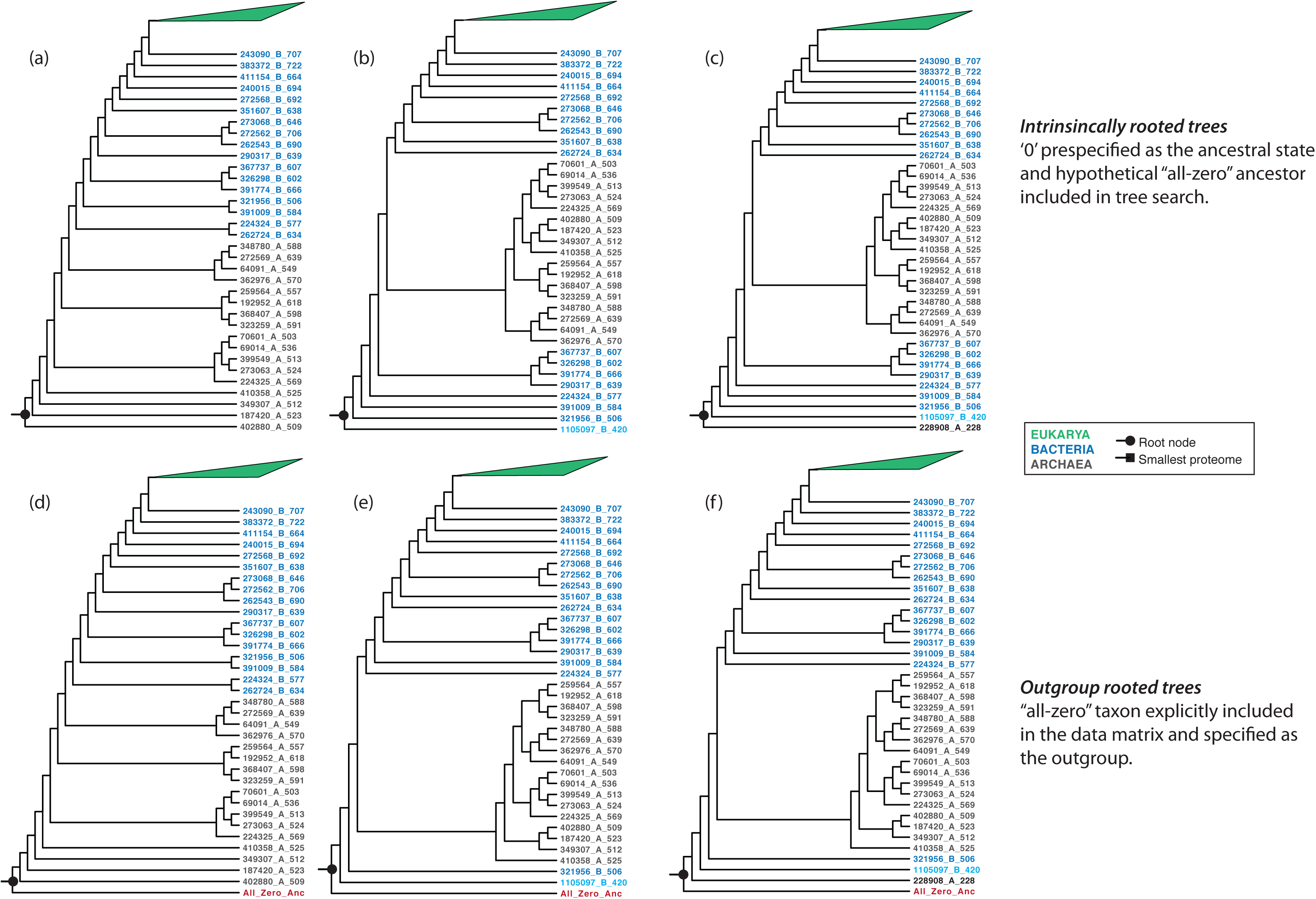
Rooting experiments (continued from Fig 1) where trees were either intrinsically rooted by including the hypothetical ancestor in tree search (a-c) or by including and explicit all-zero ancestor taxon and specifying it to be the outgroup.

**Figure S2.**
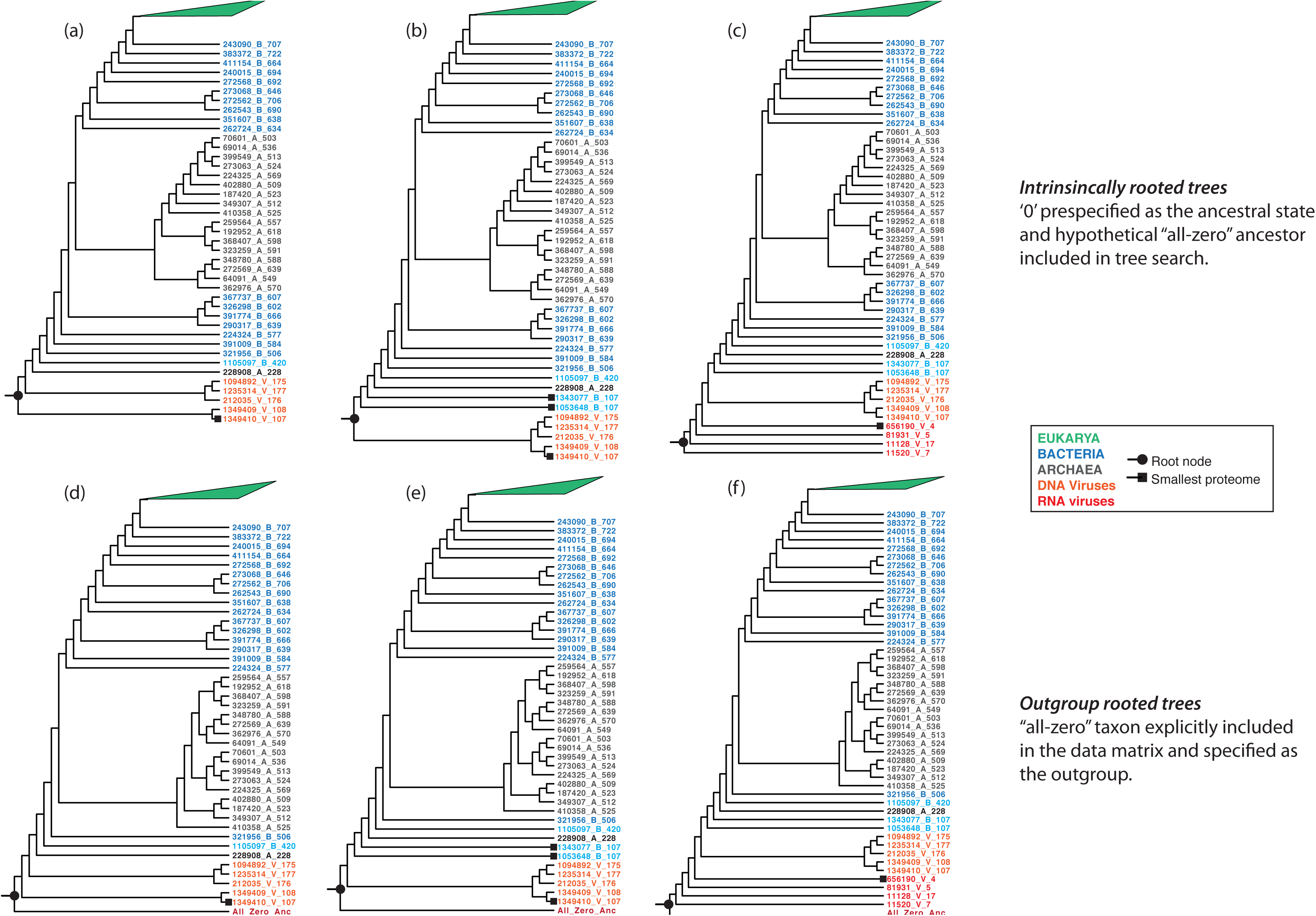
Rooting experiments (continued from Fig 2) where trees were either intrinsically rooted by including the hypothetical ancestor in tree search (a-c) or by including and explicit all-zero ancestor taxon and specifying it to be the outgroup.

**Figure S3.**
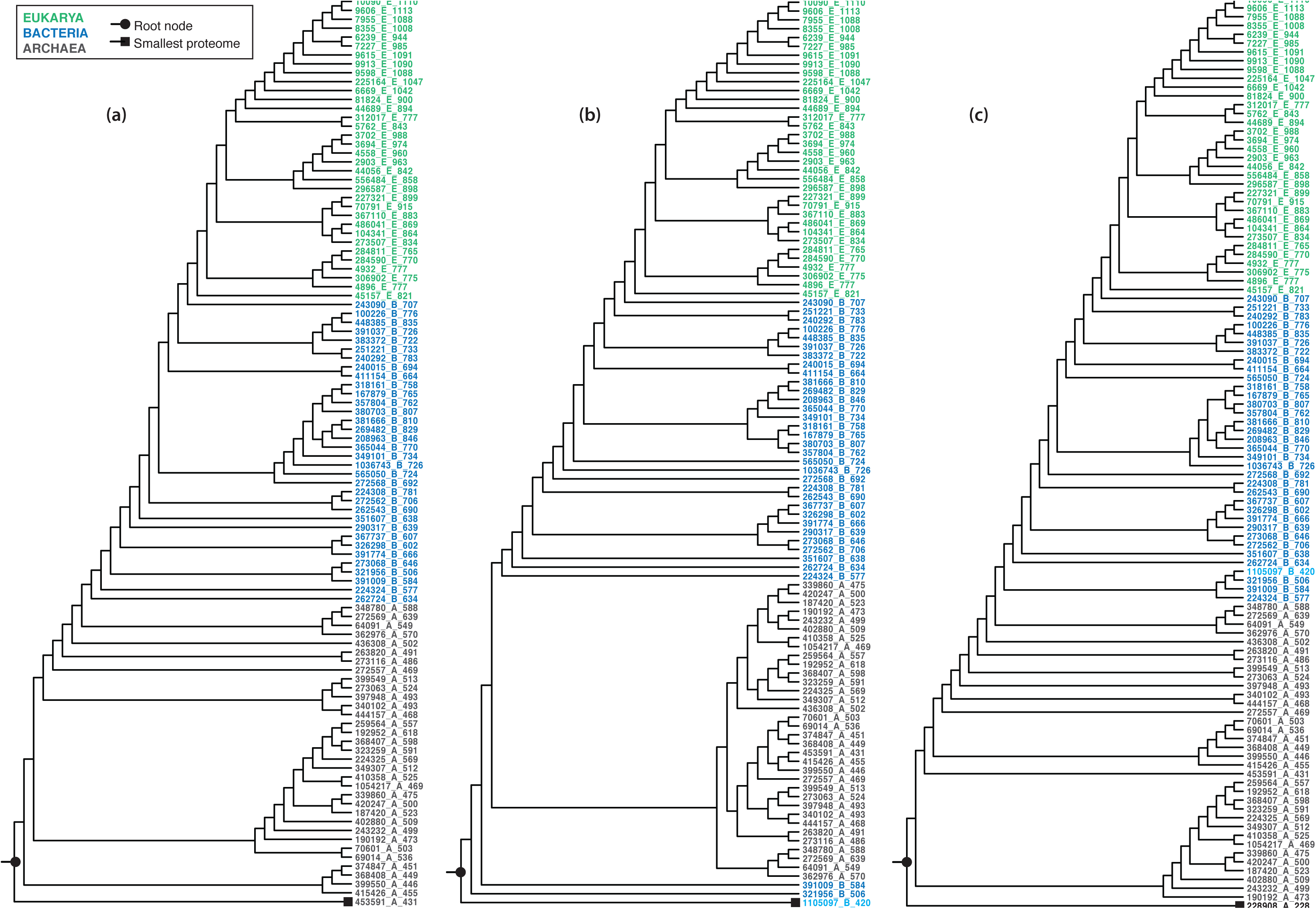
Rooting experiments (corresponding to Fig 1) with a larger taxon sampling

**Figure S4.**
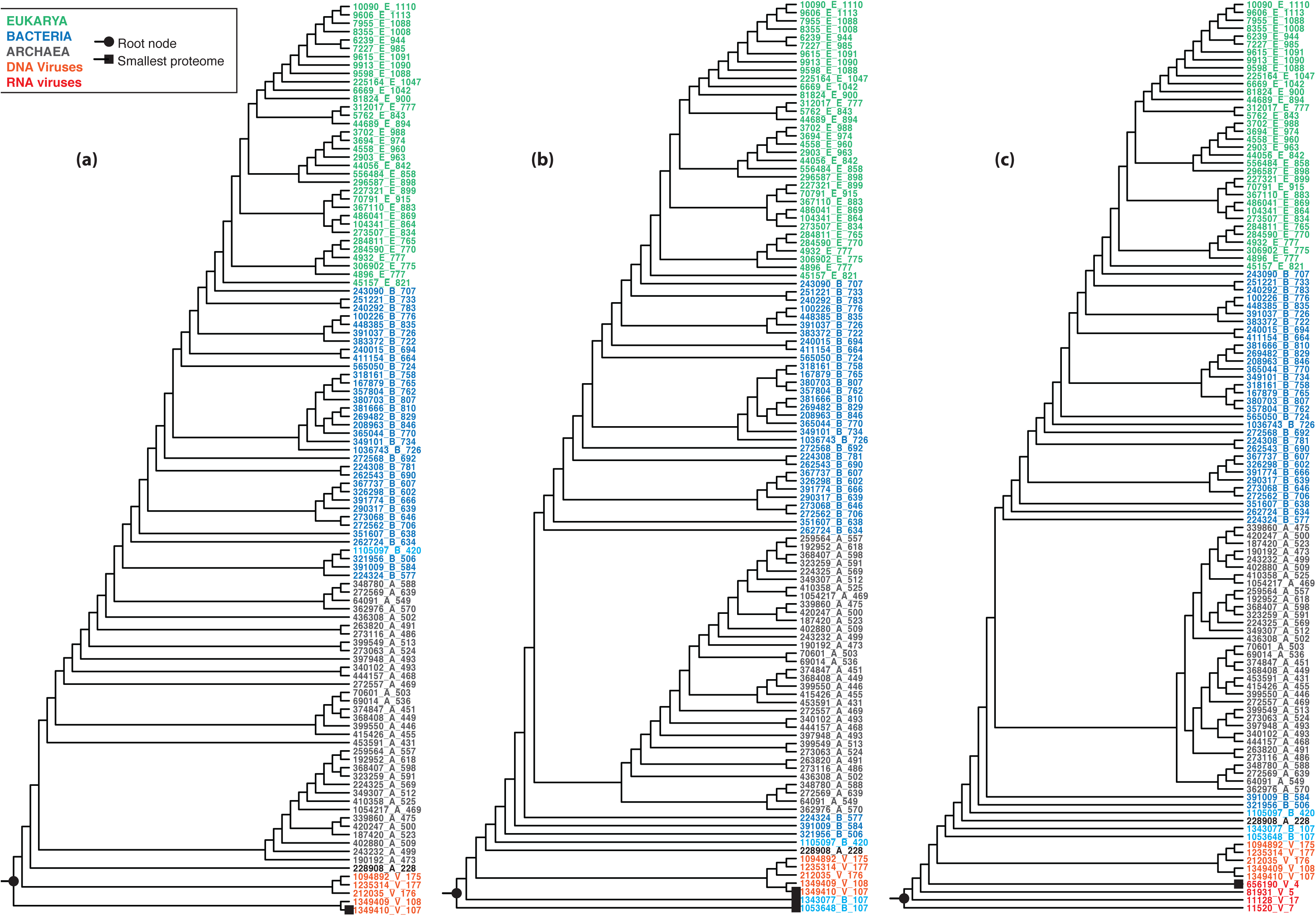
Rooting experiments (corresponding to Fig 2) with a larger taxon sampling

## References

1. Wessner, D., The origins of viruses. Nature Education, 2010. 3(9): p. 37.

2. Nasir, A. and G. Caetano-Anollés, A phylogenomic data-driven exploration of viral origins and evolution. Science Advances, 2015. 1(8).

3. Nasir, A., K. Kim, and G. Caetano-Anolles, Giant viruses coexisted with the cellular ancestors and represent a distinct supergroup along with superkingdoms Archaea, Bacteria and Eukarya. BMC Evolutionary Biology, 2012. 12(1): p. 156.

4. Murzin, A.G., et al., SCOP: A structural classification of proteins database for the investigation of sequences and structures. Journal of Molecular Biology, 1995. 247(4): p. 536–540.

5. Snel, B., P. Bork, and M.A. Huynen, Genome phylogeny based on gene content. Nature Genetics, 1999. 21(1): p. 108–110.

6. Yang, S., R.F. Doolittle, and P.E. Bourne, Phylogeny determined by protein domain content. Proceedings of the National Academy of Sciences of the United States of America, 2005. 102(2): p. 373–378.

7. Fang, H., et al., A daily-updated tree of (sequenced) life as a reference for genome research. Scientific Reports, 2013. 3.

8. Kurland, C.G. and A. Harish, Structural biology and genome evolution: An introduction. Biochimie, 2015. 119: p. 205–208.

9. Kim, K.M. and G. Caetano-Anollés, The proteomic complexity and rise of the primordial ancestor of diversified life. BMC Evolutionary Biology, 2011. 11(1).

10. Gouy, R., D. Baurain, and H. Philippe, Rooting the tree of life: the phylogenetic jury is still out. Phil. Trans. R. Soc. B, 2015. 370(1678): p. 20140329.

11. Graham, S.W., R.G. Olmstead, and S.C.H. Barrett, Rooting Phylogenetic Trees with Distant Outgroups: A Case Study from the Commelinoid Monocots. Molecular Biology and Evolution, 2002. 19(10): p. 1769–1781.

12. Morrison, D.A., Phylogenetic Analyses of Parasites in the New Millennium, in Advances in Parasitology. 2006. p. 1–124.

13. Pace, N.R., A molecular view of microbial diversity and the biosphere. Science, 1997. 276(5313): p. 734–740.

14. Wheeler, W.C., Systematics: A course of lectures. 2012: John Wiley & Sons.

15. Schwartz, R. and M. Dayhoff, Origins of prokaryotes, eukaryotes, mitochondria, and chloroplasts. Science, 1978. 199(4327): p. 395–403.

16. Woese, C.R., O. Kandler, and M.L. Wheelis, Towards a natural system of organisms: Proposal for the domains Archaea, Bacteria, and Eucarya. Proceedings of the National Academy of Sciences of the United States of America, 1990. 87(12): p. 4576–4579.

17. Swofford, D.L. and P.B. Begle, PAUP: Phylogenetic Analysis Using Parsimony: User’s Manual (Version 3.1). 1993, Champaign, IL, USA.: Illinois Natural History Survey.

18. Harish, A., A. Tunlid, and C.G. Kurland, Rooted phylogeny of the three superkingdoms. Biochimie, 2013. 95(8): p. 1593–1604.

19. Abroi, A. and J. Gough, Are viruses a source of new protein folds for organisms? - Virosphere structure space and evolution. BioEssays, 2011. 33(8): p. 626–635.

20. Abroi, A., A protein domain-based view of the virosphere-host relationship. Biochimie, 2015. 119: p. 231–243.

21. Andersson, S.G.E. and C.G. Kurland, Reductive evolution of resident genomes. Trends in Microbiology, 1998. 6(7): p. 263–268.

22. Giovannoni, S.J., J. Cameron Thrash, and B. Temperton, Implications of streamlining theory for microbial ecology. ISME J, 2014. 8(8): p. 1553–1565.

23. Dujon, B., et al., Genome evolution in yeasts. Nature, 2004. 430(6995): p. 35–44.

24. Gough, J., et al., Assignment of homology to genome sequences using a library of hidden Markov models that represent all proteins of known structure. Journal of Molecular Biology, 2001. 313(4): p. 903–919.

25. Oliveberg, M. and P.G. Wolynes, The experimental survey of protein-folding energy landscapes. Quarterly reviews of biophysics, 2005. 38(03): p. 245–288.

26. Wolynes, P.G., Evolution, energy landscapes and the paradoxes of protein folding. Biochimie, 2015. 119: p. 218–230.

27. Forterre, P., M. Krupovic, and D. Prangishvili, Cellular domains and viral lineages. Trends in Microbiology, 2014. 22(10): p. 554–558.

28. Swofford, D.L., PAUP*. Phylogenetic Analysis Using Parsimony (*and Other Methods). Version 4. 2003, Sunderland, Massachusetts: Sinauer Associates.

